# Cryptic RNA binding sites are energetically accessible and chemically addressable

**DOI:** 10.64898/2026.07.01.735868

**Authors:** Lukasz T. Olenginski, Robert T. Batey

## Abstract

Cryptic binding sites generated by local conformational dynamics have become an important concept in protein-targeted ligand discovery, yet their energetic accessibility and relevance to RNA recognition remain less well understood. Here, we use the *env8* cobalamin (Cbl) riboswitch as a model system to investigate the energetic consequences of cryptic-site formation through base displacement. Structural analysis revealed that binding of β-axial substituted Cbl derivatives displaces a conserved adenosine (A20) from the RNA core, exposing a previously hidden binding site that is subsequently occupied by the β-axial substituent. Using selective abasic substitution at this position, we quantified the energetic contributions associated with A20 in the native RNA core and with base displacement. Isothermal titration calorimetry and fluorescence measurements revealed that cryptic-site formation incurs a modest energetic penalty of ∼1.4 kcal mol^-1^. Guided by this experimentally derived framework, computational conformational sampling recapitulated cryptic-site formation in the Cbl riboswitch and identified analogous cryptic sites in structurally unrelated RNAs from HIV-1 and HCV. These cryptic-site conformers were identified within low-energy conformational windows and exposed ligand-accessible surfaces through local base displacement. Finally, a ligand previously identified to target the *env8* cryptic site bound both RNAs and yielded docking poses consistent with engagement of the newly exposed binding surfaces. Together, these results indicate that cryptic RNA binding sites can be both energetically accessible and chemically addressable, expanding the range of conformational states that may contribute to RNA ligandability.

## Main text

RNA ligandability remains challenging because few experimentally validated binding pockets are known.^1–5^ At the same time, RNA is increasingly understood as a conformational ensemble rather than a static structure.^6,7^ These observations have prompted growing interest in whether sparsely populated RNA conformations may transiently expose ligandable surfaces inaccessible in dominant structural states.^8–10^ If such conformations are energetically accessible, they could expand the repertoire of ligandable RNA sites.

Such cryptic pockets have become an important concept in protein-targeted ligand discovery, where conformational fluctuations transiently expose ligand-accessible cavities that are absent from the ground-state structure.^11–13^ Similar ideas are increasingly being explored in RNA^14^ through experimental studies of conformational heterogeneity,^9,10^ ensemble modeling,^15,16^ and computational pocket-identification^3,4,8,17^ approaches. Together, these studies suggest that transient pocket-like states may contribute to RNA ligandability. What remains less clear is the energetic cost of accessing such conformations and whether they are populated sufficiently to support productive ligand recognition.

To investigate the energetic accessibility of a cryptic RNA binding site generated by base displacement, we turned to the *env8* methylcobalamin (MeCbl)-sensing riboswitch. In our previous studies of the homologous *env2* system, we observed that β-axial substituted Cbl derivatives displace A20 from its π-stacked position between G19 and A68 within a purine spine in the core of the RNA (**Figure 1A,B**).^14^ Displacement of A20 exposes a cryptic binding site that includes the region previously occupied by the A20 nucleobase, allowing the β-axial substituent to engage this newly accessible cavity.^14^ In agreement with previous structural studies of *env8* bound to hydroxocobalamin (OHCbl) (**Figure 1B**),^18^ the *env2*-cyanocobalamin (CNCbl) structure shows A20 incorporated within the purine spine (**Figure 1C, Figure S1**).^14^ In contrast, phenyl-*p*NO_2_-Cbl binding displaces A20 from the RNA core and positions the *para*-nitrophenyl group within the cryptic binding site created by base displacement (**Figure 1C**).^14^

**Figure 1.**
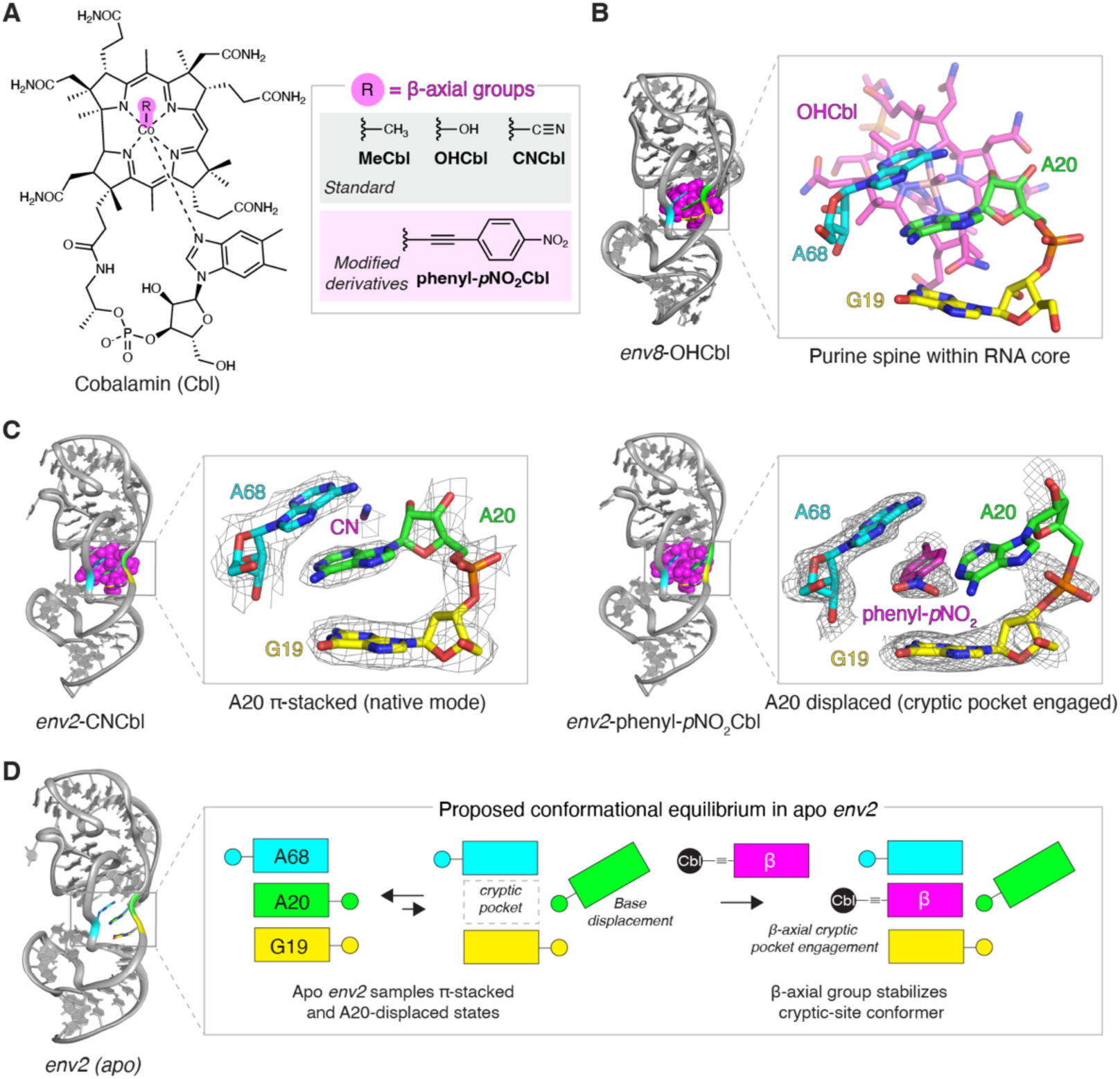
Structural basis for cryptic-site formation in the cobalamin (Cbl) riboswitch. (A) Chemical structure of Cbl highlighting the β-axial position (R). Representative native and synthetic derivatives relevant to this study are shown. (B) Structure of the *env8* riboswitch bound to OHCbl (PDB 4FRG).^18^ Expansion of the binding pocket highlights the G19-A20-A68 purine spine within the RNA core. (C) Comparison of *env2* riboswitch structures bound to CNCbl (left, PDB 9E5H) and phenyl-*p*NO_2_-Cbl (right, PDB 9E5I).^14^ In the CNCbl-bound structure, A20 is π-stacked within the G19-A20-A68 purine spine. Binding of phenyl-*p*NO_2_-Cbl displaces A20 from the RNA core, exposing a hidden binding site that is occupied by the *para*-nitrophenyl substituent. In all structural representations, the binding pocket nucleotides G19 (yellow), A20 (green), and A68 (cyan) are numbered in reference to full-length *env8* and colored; ligands (or β-axial substituents) are shown in magenta, and mesh representations correspond to a simulated annealing 2F_o_-F_c_ map where A20 and the ligand were omitted from the model are contoured at 1 σ. (D) Proposed model for cryptic-site formation in the apo riboswitch. A20 samples π-stacked and base-displaced conformations, where the latter generates a cryptic binding site. Binding of β-axial Cbl derivatives stabilizes the transient cryptic-site conformer through engagement of the newly exposed binding surface.

These structures can be interpreted as two endpoints of a local conformational equilibrium in the apo riboswitch (**Figure 1D**). In this model, A20 adopts either a π-stacked conformation within the purine spine or a displaced conformation that generates a cryptic-site conformer in which a previously hidden binding pocket becomes accessible. Binding of β-axial derivatives selectively stabilizes this cryptic-site conformer through engagement of the newly exposed binding surface. This system therefore provides a tractable model for examining the energetic consequences of base displacement and the extent to which such rearrangements can support ligand binding.

To quantify the energetic contributions associated with A20 within the native RNA core and the energetic penalty associated with A20 base displacement, we used selective abasic-site substitution at this position (**Figure 2A, Table S1**). Removal of the A20 nucleobase preserves the overall RNA scaffold while eliminating the energetic contribution of A20 to the G19-A20-A68 purine spine. Comparison of ligand binding to wild-type (WT) and abasic (Ab) *env8* aptamer domain constructs therefore provides an experimental framework for estimating both the energetic contribution associated with A20 within the RNA core and the energetic penalty associated with formation of the cryptic-site conformer. For CNCbl, which does not induce base displacement, differences in binding between WT and Ab constructs report the energetic contribution associated with A20 within the G19-A20-A68 purine spine, reflecting both stabilization of the RNA core by A20 π-stacking and minor local interactions between A20 and the CN group (**Figure 2A**). In contrast, phenyl-*p*NO_2_-Cbl binding to the Ab construct retains the favorable interactions associated with occupation of the cryptic binding site without incurring the energetic penalty associated with A20 displacement and formation of the cryptic-site conformer (**Figure 2A**).

**Figure 2.**
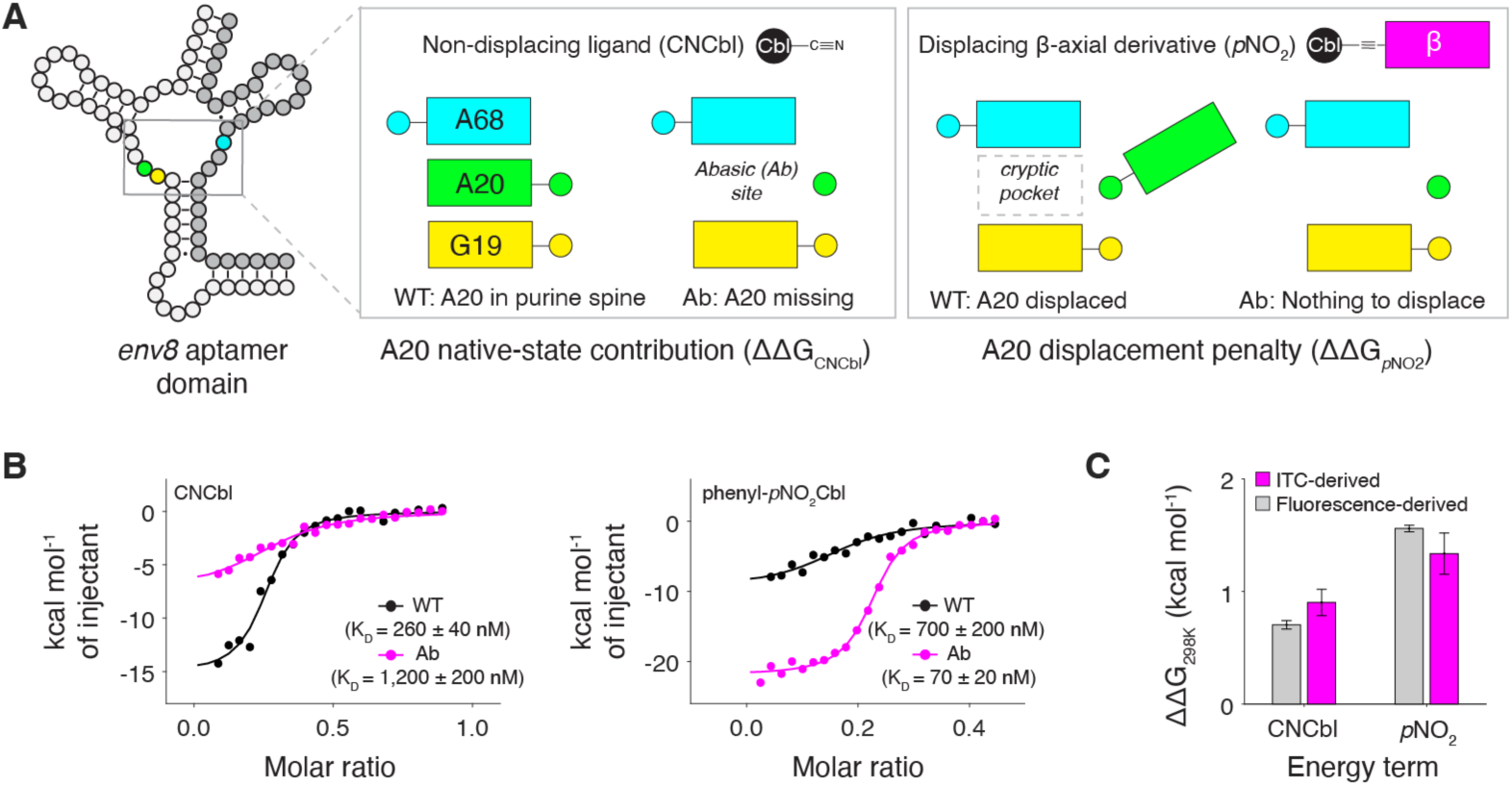
Energetic framework for cryptic-site formation through base displacement. (A) Experimental strategy used to estimate the energetic contributions associated with A20 native and cryptic conformations. Comparison of ligand binding to wild-type (WT) and abasic (Ab) *env8* constructs isolates the energetic contribution associated with A20 in the RNA core for CNCbl and the energetic penalty associated with A20 displacement for phenyl-*p*NO_2_-Cbl. (B) Representative isothermal titration calorimetry (ITC) experiments of CNCbl and phenyl-*p*NO_2_-Cbl binding to WT and Ab *env8* constructs. Representative binding isotherms and fitted dissociation constants (K_D_) from independent measurements (*n* = 3) are shown. (C) Energetic contributions derived from ITC and fluorescence experiments using PEG-linked ATTO590 derivatives of CNCbl^19^ and phenyl-*p*NO_2_-Cbl.^20^ ΔΔG values were calculated from WT-versus-Ab binding measurements and correspond to the energetic contribution of within the native RNA core (CNCbl) and the energetic penalty associated with base displacement (phenyl-*p*NO_2_-Cbl). Data are shown as mean ± SD from independent measurements (*n* = 3).

Isothermal titration calorimetry (ITC) measurements revealed that the native and cryptic conformations contribute distinct energetic terms to ligand recognition (**Figure 2B,C**). Consistent with stabilization of the native RNA core, removal of A20 in the Ab construct weakened CNCbl binding approximately five-fold relative to WT *env8*. This difference corresponds to an energetic contribution of 0.9 ± 0.1 kcal mol^-1^ associated with A20 within the purine spine, dominated by π-stacking interactions. In contrast, phenyl-*p*NO_2_-Cbl bound approximately 10-fold more tightly to the Ab construct than to the WT riboswitch. Because binding to the Ab construct does not require A20 displacement, this difference corresponds to a penalty of 1.4 ± 0.2 kcal mol⁻¹ associated with formation of the cryptic-site conformer. Notably, the energetic penalty associated with A20 displacement exceeds the energetic contribution of A20 within the native RNA core. This observation is consistent with the displacement penalty reflecting not only disruption of the G19-A20-A68 purine spine, but also additional local structural rearrangements required to generate the cryptic-site conformer. Nevertheless, occupation of the cryptic binding site by the β-axial substituent provides compensating interactions sufficient to support high affinity ligand binding.^14^

To independently evaluate these energetic estimates, we used previously developed fluorescence-based analogs of CNCbl^19^ and phenyl-*p*NO_2_-Cbl^20^ containing a PEG-linked ATTO590 fluorophore and measured binding to the corresponding WT and Ab riboswitch constructs. Despite the presence of the appended fluorophore, the fluorescence-derived ΔΔG values closely matched those obtained by ITC (**Figure 2C, Figure S2**). Estimates of 0.7 ± 0.1 kcal mol^-1^ for A20 within the native RNA core and 1.6 ± 0.1 kcal mol^-1^ for A20 displacement were in good agreement with the corresponding ITC values, providing orthogonal support for the energetic framework.

Notably, the measured base displacement penalty corresponds to only ∼2-3 RT at 298 K. In a simple two-state framework, an energetic cost of this magnitude would be expected to permit low but appreciable populations of the cryptic-site conformer (on the order of a few percent to several tens of percent). Similar energetic penalties have been reported for local RNA conformational dynamics measured by relaxation-dispersion NMR^21,22^ and chemical probing,^23^ suggesting that cryptic-site formation may be thermodynamically accessible within native RNA ensembles. These observations indicate that cryptic-site formation need not represent a rare or prohibitively costly structural excursion.

To identify analogous cryptic binding sites in unrelated RNAs, we examined experimentally determined structures containing compact purine-stacking motifs qualitatively similar to that observed in *env8*. Stem-loops from the Rev response element of HIV-1^24^ (HIV-RRE-SLII) and the internal ribosome entry site of HCV^25^ (HCV-IRES-SLIIa) were selected as representative systems based on the presence of locally π-stacked nucleobases capable of occluding potential binding surfaces (**Figure 3A,C,E**). Computational conformational searches were initiated from the experimentally determined structures and restricted to conformers within a 3 kcal mol^-1^ energy window. In each case, conformational sampling readily identified states in which the central π-stacked nucleobase adopted a displaced orientation, exposing a previously hidden binding surface within the RNA core (**Figure 3, Figure S3**).

**Figure 3.**
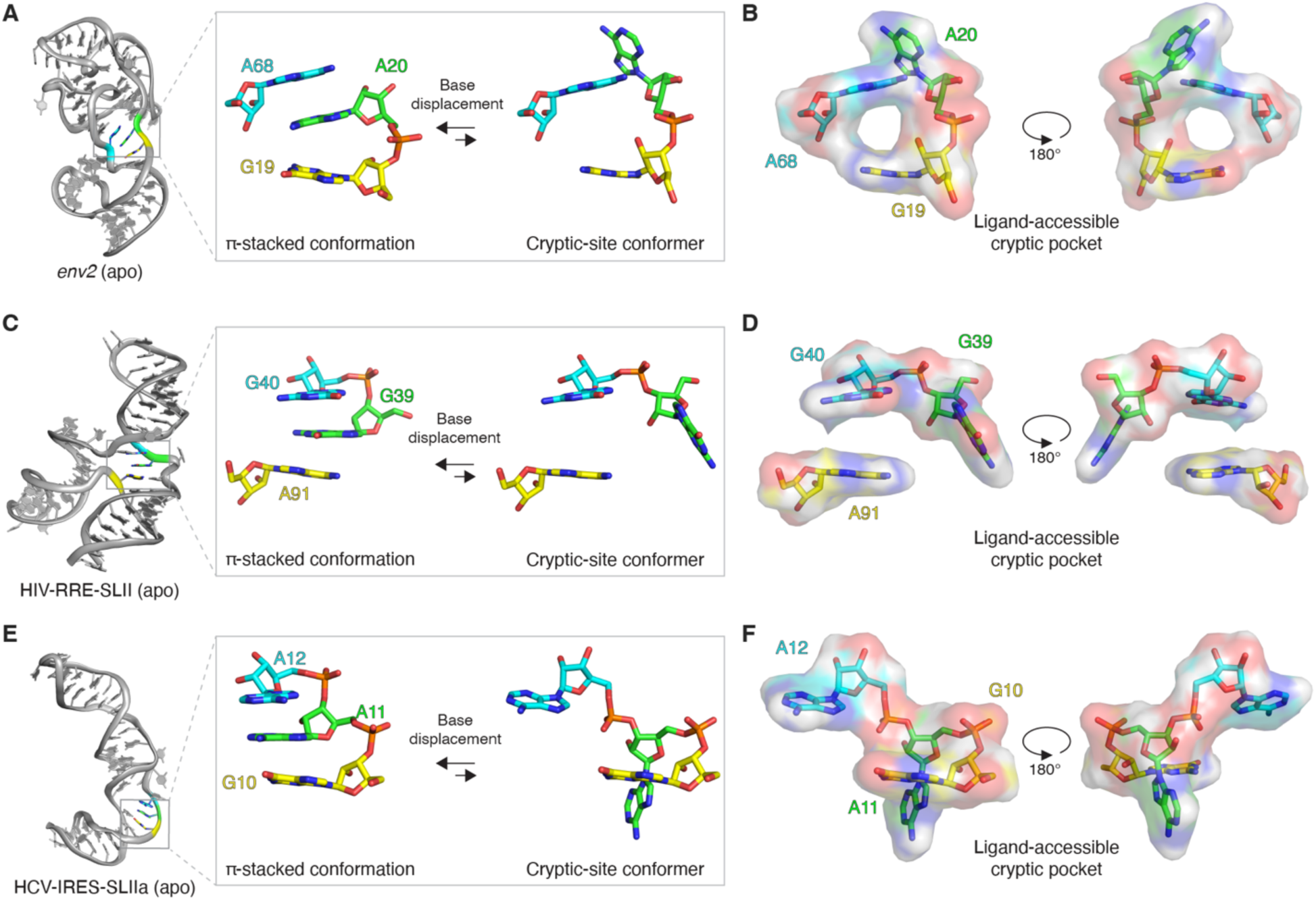
Identification of analogous cryptic binding sites in structurally unrelated RNAs. (A) Experimentally determined structure of the apo *env2* Cbl riboswitch (PDB 9MFH)^14^ highlighting the G19-A20-A68 purine spine and a representative equilibrium between the ground-state structure and a cryptic-site conformer identified through computational conformational sampling within a 3 kcal mol^-1^ energy window. The cryptic-site conformer corresponds to the lowest-energy structure in which A20 adopts a displaced conformation. (B) Surface representation of the cryptic-site conformer shown in panel A, rotated 180° about the y-axis to visualize the newly exposed cryptic binding surface generated by A20 displacement. (C) Experimentally determined structure of HIV-RRE-SLII (PDB 8UO6, chain A)^24^ highlighting the G39-G40-A91 π-stacking motif and a representative equilibrium between the ground-state structure and a cryptic-site conformer identified as in panel A. (D) Surface representation of the cryptic-site conformer shown in panel C. (E) Experimentally determined structure of HCV-IRES-SLIIa (PDB 1P5M, lowest energy NMR conformer^25^) highlighting the G10-A11-A12 π-stacking motif and a representative equilibrium between the ground-state structure and a cryptic-site conformer identified as in panel A. (F) Surface representation of the cryptic-site conformer shown in panel E.

Despite differences in global architecture, all three RNAs accessed structurally analogous cryptic-site conformers generated through local base-displacement events. In *env8*, displacement of A20 exposes the cryptic binding site (**Figure 3A,B**) occupied by β-axial Cbl substituents.^14^ Similar rearrangements were observed in HIV-RRE-SLII, where displacement of G39 from the G39-G40-A91 π-stacking motif generated a ligand-accessible cavity (**Figure 3C,D, Figure S4**), and in HCV-IRES-SLIIa, where displacement of A11 from the G10-A11-A12 motif exposed an analogous binding surface (**Figure 3E,F**). In each system, the displaced nucleobase vacated a region of the RNA core that becomes accessible to potential ligand engagement.

While the energetic values obtained from these computational conformational searches should be interpreted cautiously and do not establish the populations of the resulting conformers, the identification of cryptic-site conformers within low-energy conformational windows is consistent with the experimentally derived framework established for *env8*. Together, these results suggest that cryptic-site formation through local base displacement is not unique to the *env8* riboswitch. Instead, similar pocket-forming rearrangements can be accessed in structurally unrelated RNAs through modest local conformational changes, suggesting that cryptic binding surfaces may represent a broader feature of RNA conformational landscapes.

To evaluate whether these newly identified cryptic binding sites could support small molecule recognition, we leveraged a focused ligand set previously developed against the *env8* cryptic site.^14^ In earlier work, computational screening of an RNA-focused library identified eight candidate ligands (**E1**-**E8**) containing biphenyl-like scaffolds capable of engaging the cryptic binding site formed upon A20 displacement (**Figure 4A**).^14^ Experimental testing revealed that two compounds, **E2** and **E5**, bound the *env8* riboswitch with 30-50 μM affinity and competitive behavior consistent with engagement of the Cbl-binding pocket.^14^

**Figure 4.**
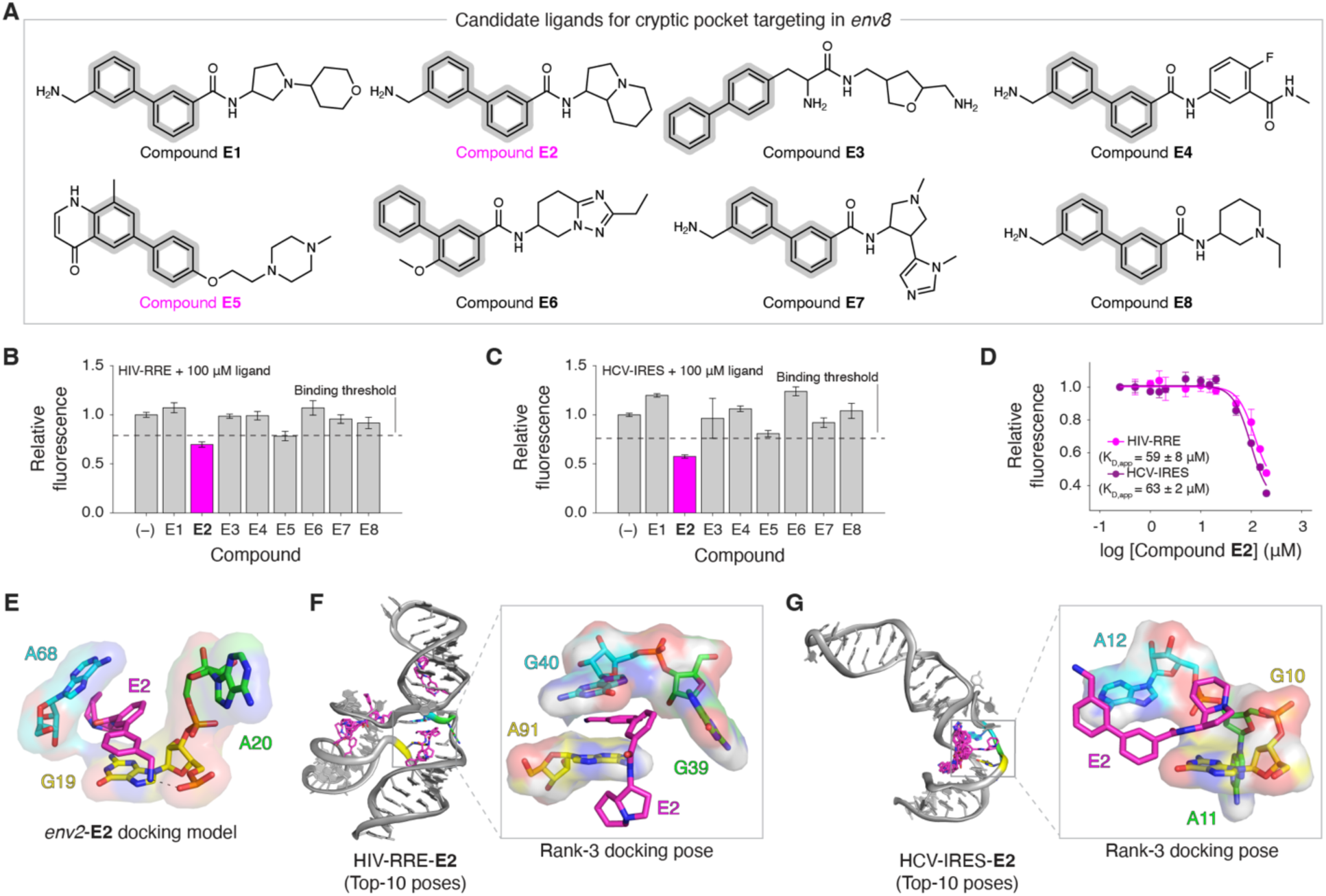
Chemical addressability of cryptic RNA binding sites. (A) Chemical structures of compounds **E1**-**E8** with biphenyl-like scaffolds identified from computational screening of an RNA-focused library for compounds capable of engaging the *env8* cryptic binding site. Ligands **E2** and **E5**, previously shown to bind the *env8* riboswitch, are highlighted in magenta. (B) Thiazole orange (TO) displacement assay of ligands **E1**-**E8** against HIV-RRE-SLII. Dashed line indicates the binding threshold used to identify candidate ligands for follow-up analysis. (C) TO displacement data of ligands **E1**-**E8** against HCV-IRES-SLIIa as in panel B. (D) Representative TO displacement titrations of E2 against HIV-RRE-SLII and HCV-IRES-SLIIa. Solid lines represent fits to the binding model used to estimate apparent K_D_ (K_D,app_) values. Data in panels B-D are shown as mean ± SD from independent measurements (*n* = 3). (E) Top-ranked docking pose of **E2** within the *env8* cryptic binding site. Consistent with previous competitive binding studies, the biphenyl scaffold is positioned within the cryptic site exposed by A20 displacement. (F) Top 10 docking poses obtained for **E2** against the cryptic-site conformer of HIV-RRE-SLII. The inset highlights the third-ranked (Rank-3) docking pose, where the biphenyl moiety occupies the newly exposed cryptic site. (G) Top 10 docking poses obtained for **E2** against the cryptic-site conformer of HCV-IRES-SLIIa. The inset highlights the Rank-3 pose, where the aliphatic, fused 5,6-ring system extends into the newly exposed pocket. Computational docking was performed using unbiased ligand placement and conformational ligand sampling.

The same ligand panel was subsequently evaluated against HIV-RRE-SLII and HCV-IRES-SLIIa using a thiazole orange (TO) displacement assay,^14,26–28^ where attenuation of TO fluorescence is indicative of ligand binding. Among the eight compounds tested, **E2** was the only compound that reproducibly reduced TO fluorescence in a manner consistent with binding both RNAs (**Figure 4B,C**). Follow-up titrations yielded apparent K_D_ (K_D,app_) values of 59 ± 8 μM for HIV-RRE-SLII and 63 ± 2 μM for HCV-IRES-SLIIa (**Figure 4D**), placing all three RNAs within a similar affinity regime. Thus, a ligand originally identified through targeting of the *env8* cryptic site retained comparable binding activity against two structurally distinct RNAs containing analogous cryptic sites.

Previous structural and competitive binding studies of *env8* support a binding mode in which the biphenyl scaffold of **E2** occupies the cryptic binding site exposed by A20 displacement (**Figure 4E**).^14^ We therefore asked whether **E2** could similarly engage these sites in HIV-RRE-SLII and HCV-IRES-SLIIa (**Figure 3).** Unbiased ligand docking against the cryptic-site conformers of both RNAs yielded favorable poses consistent with cryptic-site engagement (**Figure 4F,G**). In HIV-RRE-SLII, the biphenyl moiety occupied the newly exposed cryptic site in a manner analogous to that observed in *env8* (**Figure 4F, Figure S5A**). In contrast, docking to HCV-IRES-SLIIa positioned the biphenyl scaffold adjacent to the cryptic site, allowing the aliphatic, fused 5,6-ring system of **E2** to extend into the newly exposed pocket (**Figure 4G, Figure S5B**). While our docking analysis does not establish the precise binding mode of **E2**, it provides a structural hypothesis for the observed binding activity.

Together, these results demonstrate that cryptic RNA binding sites can be both energetically accessible and chemically addressable. Using the *env8* Cbl riboswitch as a model system, we show that cryptic-site formation through base displacement incurs only a modest energetic penalty and can support productive small molecule recognition. Guided by this experimentally derived framework, we identified analogous cryptic sites in structurally unrelated RNAs containing similar local π-stacking motifs. In each case, displacement of a central nucleobase exposed a hidden ligand-accessible surface, suggesting that cryptic-site formation may arise naturally from this structural feature. Similar π-stacking networks are common features of structured RNAs, where they contribute to folding stability and long-range communication.^29^ Our findings raise the possibility that these motifs may also provide productive starting points for identifying cryptic RNA binding sites.

More broadly, these observations suggest that ligandable RNA space may extend beyond binding surfaces evident in static structures. If recurrent π-stacking motifs can access cryptic-site conformers through energetically accessible local conformational dynamics, then similar motifs may provide a structural framework for identifying previously unrecognized RNA binding sites. The emergence of computational approaches for identifying ligandable and cryptic RNA pockets from structural data further suggests that experimentally grounded frameworks describing the energetic accessibility and structural origins of such sites may aid in their systematic identification across broader classes of RNA.^17^ More generally, RNA regions containing π-stacked nucleobases contributing to local structural organization but are not rigidly constrained by extensive secondary or tertiary interactions may be particularly susceptible to the conformational fluctuations that give rise to cryptic binding pockets. In this view, cryptic-site formation is not simply a consequence of RNA dynamics, but a potentially exploitable feature of RNA structure that expands the range of conformations available for small molecule recognition and therapeutic targeting.

## Author Information

### Author Contributions

L.T.O. and R.T.B. designed the research. L.T.O. collected and analyzed all experimental and computational data and wrote the paper with feedback from all authors. R.T.B. supervised the project.

### Notes

The authors declare the following competing financial interest(s): R.T.B. serves on the Scientific Advisory Boards of MeiraGTx and Mol Horizon. R.T.B. and L.T.O. hold equity stakes in Mol Horizon.

## Acknowledgements

We thank the Shared Instruments Pool (RRID: SCR_018986) of the Department of Biochemistry at the University of Colorado Boulder for the use of the MicroCal ITC200 (S10RR026516) instrument. We acknowledge A. Erbse for her assistance with the resources utilized at the University of Colorado Boulder.

## Abbreviations

Cbl: cobalamin;
OHCbl: hydroxocobalamin;
CNCbl: cyanocobalamin;
WT: wild-type;
Ab: abasic;
ITC: isothermal titration calorimetry;
RRE: Rev response element;
IRES: internal ribosomal entry site;
TO: thiazole orange;
K_D_: dissociation constant;
K_D,app_: apparent K_D_.

