## Supplemental Information for "Cryptic RNA binding sites are energetically accessible and chemically addressable"

**This file includes:**

| | |
|---|---|
| Materials and methods | 2-3 |
| Table S1 | 4 |
| Figures S1-S5 | 5-9 |
| References | 10 |

### Materials and methods

#### RNA preparation

For abasic RNA experiments, single-stranded RNA oligonucleotides (**Table S1**) were purchased from Dharmacon (Revvity). Wild type (WT) and abasic (Ab) *env8* aptamer domain constructs were annealed by equimolar mixing of 5′- and 3′-ends (1 mM) in Milli-Q $H_2O$, incubating for 30 minutes at 37 °C, and equilibration to room temperature. Stem-loops from the Rev response element of HIV-1 (HIV-RRE-SLII[5]) and the internal ribosome entry site of HCV (HCV-IRES-SLIIa[6]) (**Table S1**) were prepared using DNA templates amplified by PCR, transcribed with T7 RNA polymerase, and purified using preparative denaturing polyacrylamide gel electrophoresis.[1] Purified RNA was buffer exchanged and concentrated into Milli-Q $H_2O$ using centrifugal concentrators (Millipore-Sigma). Final RNA concentrations were calculated using absorbance at 260 nm and extinction coefficients determined from the summation of the individual bases. Prior to all binding experiments, RNA was heated at 95 °C for 2 minutes, incubated on ice for 10 minutes, and then allowed to equilibrate to room temperature.

#### Isothermal titration calorimetry (ITC)

RNA was exchanged into a 1X RNA binding buffer (50 mM 4-(2-hydroxyethyl)-1-piperazineethanesulfonic acid (HEPES) pH 8.0, 100 mM KCl, 10 mM NaCl, and 1 mM $MgCl_2$) by successive washing in centrifugal concentrators (Millipore-Sigma). RNA was diluted up to a final concentration of 20 µM and titrated with ligand (dissolved in the flow-through of the final wash from buffer exchange) at 50-100 µM. All titrations were performed at 25 °C using a MicroCal ITC200. Data were fit to a single-site binding model using Origin 7 ITC software (MicroCal). Energetic contributions associated with A20 π-stacking and base displacement were calculated from WT-versus-Ab binding measurements using $\Delta\Delta G = RT \ln(K_{D,2}/K_{D,1})$.

#### Cobalamin (Cbl) fluorescence binding assays

For fluorescence induction titrations, 60 µL reactions containing 100 nM CNCbl-5xPEG-ATTO590[2] or phenyl-*p*$NO_2$Cbl-5xPEG-ATTO590,[3] 1X RNA binding buffer supplemented with 0.01% Nonidet P-40, and increasing amounts of WT or Ab *env8* constructs were incubated for 1 hour at room temperature. From each reaction, 50 µL was transferred to a 384-well plate (Corning) and ATTO590 fluorescence was monitored (594 nm excitation, 620-670 nm emission) at room temperature using a CLARIOstar Plus microplate reader (BMG Labtech). Fluorescence values were integrated across the emission spectrum and normalized to reactions lacking RNA. Binding data were fit using a quadratic binding model in MATLAB (v2024b) assuming a probe concentration of 100 nM. Because some RNA-probe interactions did not fully saturate within the experimental concentration range, data were analyzed using multiple endpoint-constrained fitting strategies in which the maximum fluorescence response ($F_{max}$) was fixed or allowed to vary within defined bounds. Fluorescence-derived energetic terms were calculated from WT-versus-Ab binding measurements and compared across fitting strategies, with the intermediate $F_{max}$-constrained model (0.8-1.4) used for comparison to ITC-derived energetic parameters.

**Computational conformational sampling**

For putative cryptic pocket modeling, apo structures of the *env2* Cbl riboswitch (PDB 9MFH[4]), HIV-RRE-SLII (PDB 8UO6[5], focusing on chain A), and HCV-IRES-SLIIa (1P5M[6], the lowest energy NMR conformer) were imported into Molecular Operating Environment (MOE, v2022.02, Chemical Computing Group) and prepared using the Quick Prep protocol with default parameters. To permit local conformational sampling within each target π-stacking motif, all atoms belonging to unpaired nucleotides within the motif were designated as flexible, while the remainder of the RNA was held fixed. Conformational searches were then performed using default MOE parameters with modification to the rejection (100) and iteration (500) limits as well as the MM iteration (200) and restricted to structures within a 3 kcal $mol^{-1}$ energy window. The resulting ensembles consisted of conformers generated through local sampling of the flexible nucleotides together with their corresponding calculated energies. Representative cryptic-site conformers were selected as low energy structures in which the central nucleobase of the stacking motif adopted a displaced conformation. Similar base-displaced states were observed throughout the resulting conformational ensembles, yielding comparable cryptic binding surfaces.

**Thiazole orange (TO) binding assay**

Ligands previously developed against the *env8* cryptic site[4] were screened against HIV-RRE-SLII and HCV-IRES-SLIIa. Enamine catalog identifiers are Z4539256213 (**E1**), Z3515426302 (**E2**), Z4144805645 (**E3**), Z8153471773 (**E4**), Z2466243364 (**E5**), Z1209546900 (**E6**), Z7621749448 (**E7**), and Z4116044018 (**E8**). For screening, thiazole orange (TO) displacement assays were performed using 60 µL reactions containing 0.5 µM RNA, 1 µM TO, and 100 µM competing ligand in 1X RNA binding buffer. Fluorescence values (F) were normalized to reactions lacking ligand. TO displacement was interpreted using a one-site competition model and RNA-specific fluorescence thresholds (F*) corresponding to an apparent $K_D$ ($K_{D,app}$) threshold of 200 µM were calculated. Ligands were classified as binders when $F \leq F^*$, corresponding to $K_{D,app} \leq 200$ µM under the model assumptions. For follow-up binding measurements, TO displacement titrations were carried out using increasing ligand concentrations under otherwise identical conditions. Data were fit to a four-parameter logistic (sigmoidal dose-response) model in MATLAB (v2024b). $K_{D,app}$ values were calculated from $IC_{50}$ values using the measured TO dissociation constants for HIV-RRE-SLII (1.1 µM) and HCV-IRES-SLIIa (1.8 µM).[4]

**Computational docking**

Representative cryptic-site conformers (from the computational conformational searches described above) of the *env2* Cbl riboswitch, HIV-RRE-SLII, and HCV-IRES-SLIIa apo RNAs were imported into MOE and prepared for docking using the Quick Prep protocol with default parameters. The SMILES string for ligand **E2** was imported into MOE, assigned the dominant protonation state at pH 7, and energy-minimized using default parameters. Unbiased rigid-receptor docking was performed without restricting **E2** placement, with 30 ligand poses generated and the top 10 ranked by docking score retained for analysis.

**Table S1.** RNA constructs used in this study.

| Name | Sequence (5′-to-3′) |
|---|---|
| *env8* (5′-strand, WT) | GGC CUA AAA GCG UAG UGG GAA AGU GAC GUG AAA UUC GUC CAG AUC GCC |
| *env8* (5′-strand, Ab) | GGC CUA AAA GCG UAG UGG GrabA AGU GAC GUG AAA UUC GUC CAG AUC GCC |
| *env8* (3′-strand) | GGC GAU ACG GUU AUA CUC CGA AUG CCA CCU AGG CC |
| HIV-RRE-SLII | GGC ACU AUG GGC GCA GUG UCA AUG GAC GCU GAC GGU ACA GGC CAG ACA AUU AUU GUC UGG UAU AGU GCC |
| HCV-IRES-SLIIa | GGC UGU GAG GAA CUA CUG UCU UCA CGC CUU CGG GAG UGU CGU GCA GCC UCC AGC C |

a. The *env8* wild-type (WT) and abasic (Ab) aptamer domain constructs were constructed by annealing their respective 5′-strands to the common 3′-strand. Here, "rab" refers to an abasic RNA nucleotide that is incorporated at position A20 (using full-length numbering).
b. HIV-RRE-SLII is the HIV Rev-response element (RRE) stem-loop II NMR construct from Choi and co-workers.[5]
c. HCV-IRES-SLIIa is the HCV internal ribosome entry site (IRES) stem-loop IIa NMR construct from Puglisi and co-workers.[6]

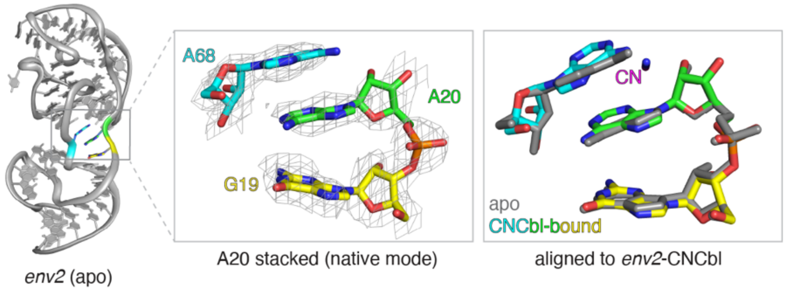


**Figure S1.** Comparison of apo and CNCbl-bound conformations of the *env2* Cbl riboswitch. (Left) Experimentally determined structure of the apo *env2* riboswitch (PDB 9MFH[4]) highlighting the G19-A20-A68 purine spine. (Middle) Expanded view of the purine spine in the apo structure, showing A20 in the π-stacked conformation. (Right) Superposition of the apo (gray) and CNCbl-bound (colored, PDB 9E5H[4]) structures aligned using the purine spine nucleotides. The close correspondence between the two structures indicates that the CNCbl-bound state adopts the native π-stacked conformation of the purine spine. In all structural representations, the binding pocket nucleotides G19 (yellow), A20 (green), and A68 (cyan) are numbered in reference to full-length *env8* and colored; the β-axial group is shown in magenta, and mesh representations correspond to a simulated annealing $2F_o$-$F_c$ map where A20 and the ligand (if present) were omitted from the model are contoured at 1 σ.

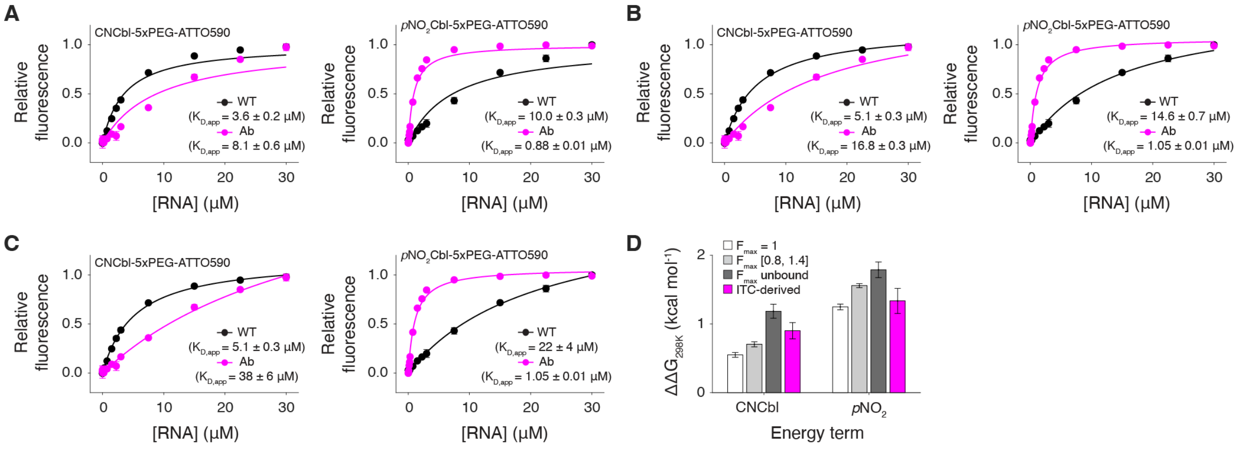


**Figure S2.** Fluorescence-derived energetic contributions of A20 contributions within the RNA core and base displacement. (A) Fluorescence induction titrations of CNCbl-5xPEG-ATTO590[2] and phenyl-*p*$NO_2$-Cbl-5xPEG-ATTO590[3] against WT and Ab *env8* constructs using a fixed endpoint model ($F_{max}$ = 1). Apparent dissociation constants ($K_{D,app}$) were obtained by fitting the data to a quadratic binding model. (B) Fluorescence induction titrations analyzed using a constrained endpoint model in which $F_{max}$ was allowed to vary between 0.8 and 1.4. This fitting strategy was used for comparison to ITC-derived energetic parameters in the main text. (C) Fluorescence induction titrations analyzed using an unconstrained endpoint model in which $F_{max}$ was allowed to vary freely. (D) Energetic contributions associated with A20 contributions within the purine spine (CNCbl) and base displacement (phenyl-*p*$NO_2$-Cbl) calculated from WT-versus-Ab binding measurements for each fitting strategy and compared to ITC-derived values. Data are shown as mean ± SD from independent measurements (*n* = 3).

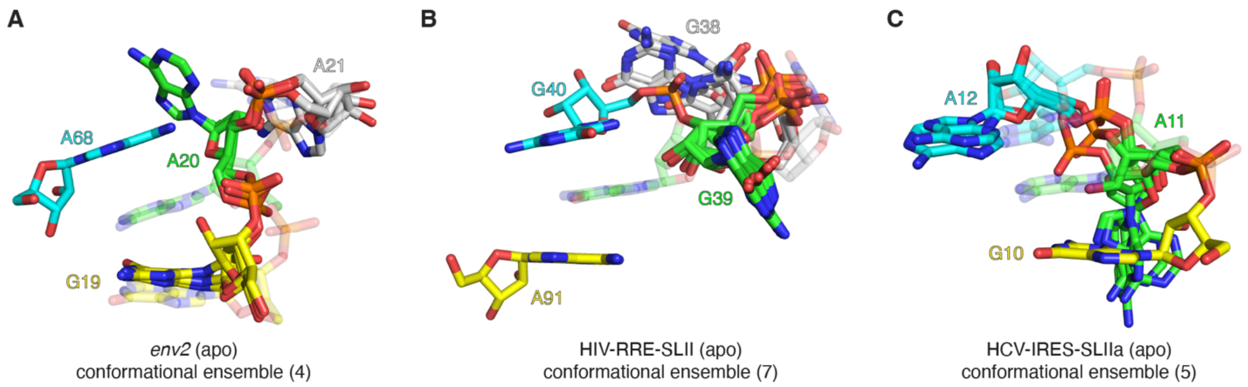


**Figure S3.** Conformational ensembles generated during putative cryptic-site modeling. (A) Conformational ensemble generated from the apo *env2* riboswitch structure during local conformational sampling of the G19-A20-A68 purine spine within the 3 kcal $mol^{-1}$ energy window. (B) Conformational ensemble generated from the apo HIV-RRE-SLII structure during local conformational sampling of the G39-G40-A91 π-stacking motif. (C) Conformational ensemble generated from the apo HCV-IRES-SLIIa structure during local conformational sampling of the G10-A11-A12 π-stacking motif. In all structural representations, the experimentally determined structure is shown as transparent sticks and the resulting ensemble of conformers are shown as opaque sticks. Numbers in parentheses indicate the total number of conformers identified within the 3 kcal $mol^{-1}$ energy window. Nucleobases comprising the target π-stacking motifs are colored as in the main text, while adjacent nucleotides included as flexible elements of the conformational search are shown in white and colored by atom type

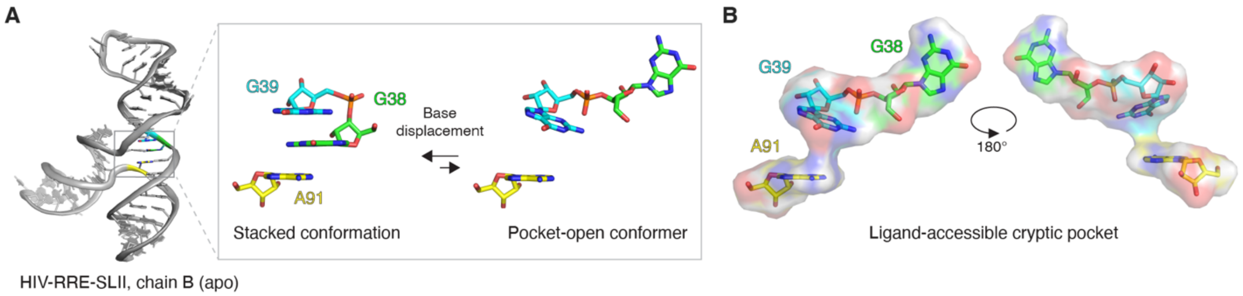


**Figure S4.** Identification of a cryptic-site conformer in HIV-RRE-SLII. (A) Experimentally determined structure of HIV-RRE-SLII (PDB 8UO6, chain B[5]) highlighting the G38-G39-A91 π-stacking motif and a representative equilibrium between the ground-state structure and a cryptic-site conformer identified through computational conformational sampling. The cryptic-site conformer corresponds to a low-energy structure in which G38 adopts a displaced conformation. (B) Surface representation of the cryptic-site conformer shown in panel A, rotated 180° about the y-axis to visualize the newly exposed cryptic binding surface generated by G38 displacement. Despite differences in the local π-stacking arrangements observed between chains A and B, base-displaced cryptic-site conformers were identified in both structures.

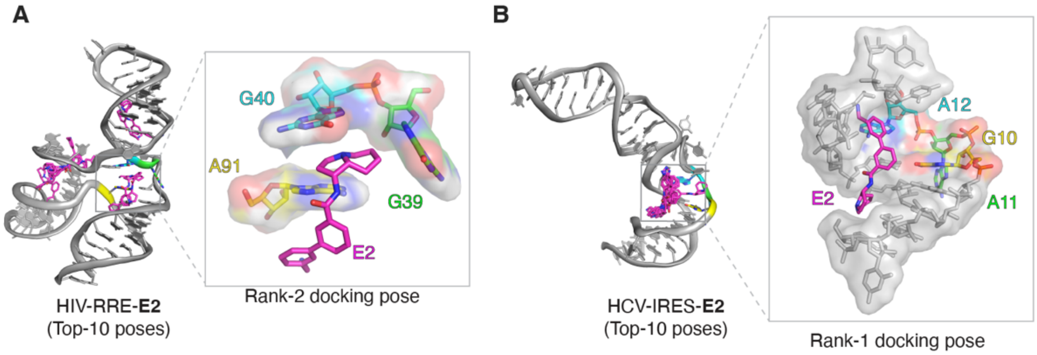


**Figure S5.** Distribution of docking poses for **E2** in cryptic-site conformers of HIV-RRE-SLII and HCV-IRES-SLIIa. (A) Top 10 docking poses obtained for **E2** against the cryptic-site conformer of HIV-RRE-SLII. The inset highlights the second-ranked (Rank-2) docking pose, in which the fused 5,6-ring system occupies the cryptic binding site while the biphenyl scaffold extends along the adjacent RNA surface. (B) Top 10 docking poses obtained for E2 against the cryptic-site conformer of HCV-IRES-SLIIa. The inset highlights the top-ranked (Rank-1) docking pose, in which the ligand remains positioned adjacent to the pocket. Computational docking was performed as described in the Methods using unbiased ligand placement and conformational ligand sampling.
